# Ghrelin and Mboat4 are lost in Serpentes

**DOI:** 10.1101/2025.05.18.653663

**Authors:** Rui Pinto, Raquel Ruivo, Josefin Stiller, Diogo Oliveira, L. Filipe C. Castro, Rute da Fonseca

## Abstract

Exploring the evolution of gene networks associated with metabolic/energetic homeostasis can yield key insights into the adaptive landscapes governing the physioology of extant lineages. Here, we investigate a key hormonal module of energy metabolism in reptiles. Ghrelin (*GHRL*), also known as the “*hunger hormone*”, is a multifunctional gastric peptide, involved in appetite, food intake and body weight regulation. We examined the genomes of 112 species comprising members of the Squamata, Testudines, Crocodilia and Rhynchocephalia and provide ample evidence that *GHRL* was independently lost in snakes, chameleons and toadhead agamas. In accordance, the enzyme responsible for ghrelin acylation and essential for its activity (membrane bound O-acyltransferase domain containing 4), is also eroded in these lineages. We suggest that the loss of this hormonal signalling system parallels critical modifications in energy metabolism, including a lower need to stimulate fatty acid oxidation during fasting in the locomotor muscles of snakes compared to other groups.

## 1. Introduction

Digestive physiology is intertwined with a species’ feeding ecology, diet and energy expenditure [1–6]. Many reptiles exhibit extreme feeding behaviours, including the ability to endure prolonged periods of fasting that can last for months or even more than a year [7,8]. Long fasting times have led to adjustments that reduce the metabolic costs in between feeding and digestion periods [9,10]. Likewise, the ingestion of large meals after fasting led to parallel digestive modifications, including increases in gastric acid secretion [11] and metabolic rate related to digestion – the latter named specific-dynamic action or SDA response (figure 1a). While accounting for a small portion of an individual’s daily energy budget in birds and endothermic mammals, the SDA response is rather energy-expensive in ectothermic sauropsids, amphibians and fish [12–14]. In order to accommodate large meals and long digestive times, reptiles also present unique adaptations regarding gastrointestinal morphology and physiology [15–19]; [20– 22]. At the genetic level, changes in gene expression upon feeding are also observed in snakes and positive selection has been determined in metabolism-related genes (e.g. *ADD1, VAPB, SREBF2*) [23]. Yet, whether the extreme physiological changes associated with fasting and digestion implicate more notable alterations in gene repertoire (e.g. gene duplication and loss) is unknown [24,25].

**Figure 1.**
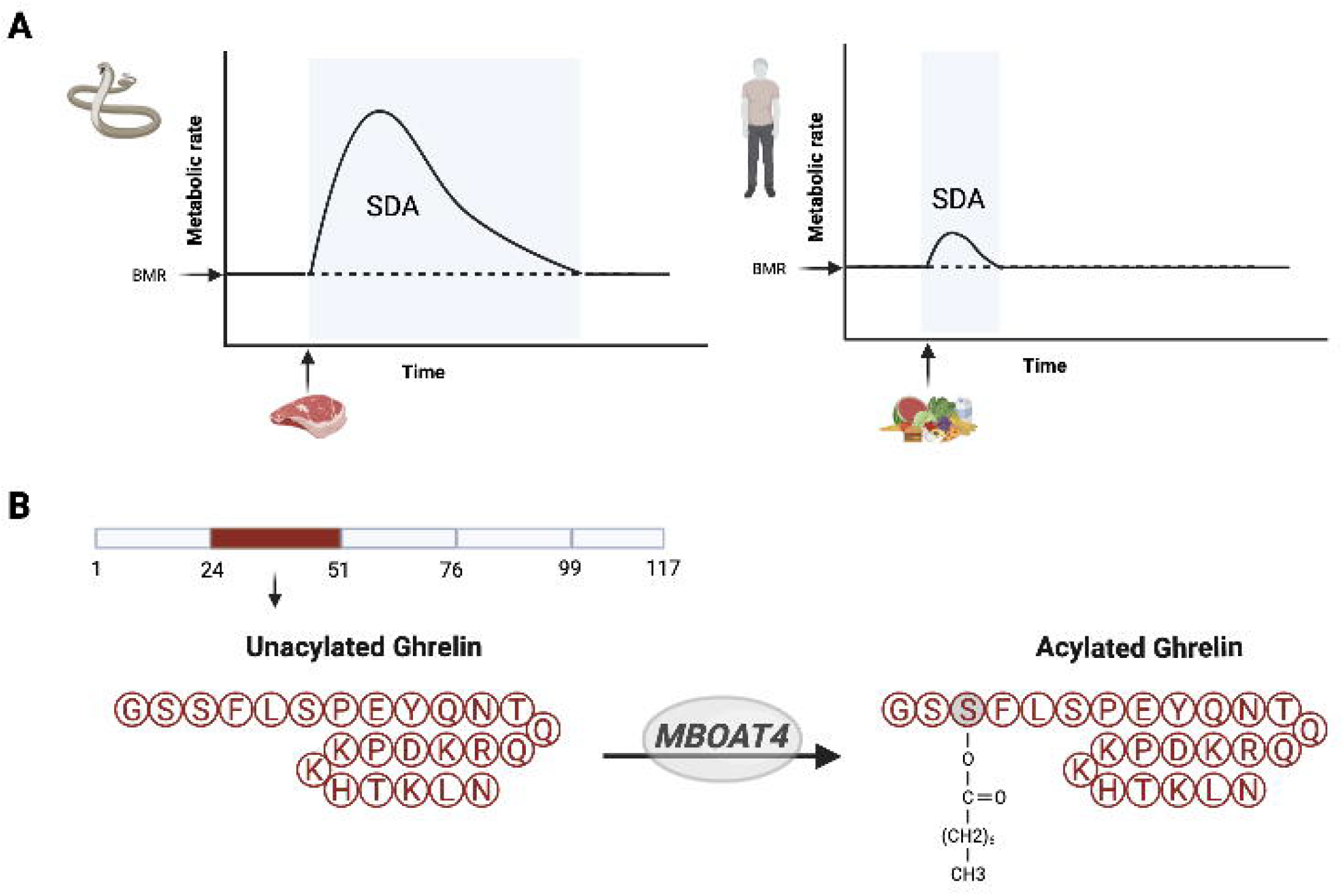
Ghrelin system, as a master regulator of energy homeostasis, is interconnected with the postprandial response. (*a*) Species with longer time between meals, like sit-and-wait foraging snakes, typically have larger meals relative to body weight, which elicit a more intense and highly energetically-costly metabolic response than that observed in mammals, birds and fishes *[12–14]*. (*b*) Preproghrelin is cleaved into unacylated GHRL and is acylated through esterification of a fatty acid on the third aminoacid (Serine 3) by an enzyme encoded by the gene *Mboat4*.

Here, we assess the functional coding status of the hormone ghrelin (*GHRL*) across 112 species of non-avian diapsids. *GHRL* regulates appetite and metabolism based on fed/fasting states (e.g. [26]), and exists as two functional states (figure 1b). The conversion of the unacylated *GHRL* (UnG) into acylated *GHRL* (AcG) is catalysed by the Membrane Bound O-Acyltransferase Domain Containing 4 enzyme (*MBOAT4*) [27]. In mammals, UnG levels increase during fasting [28], while *MBOAT4* levels decrease and AcG levels remain unchanged [27]. Feeding increases the expression of *MBOAT4* and subsequently the acylation of UnG to AcG [27]. AcG then binds to growth hormone secretagogue receptor-1a (*GHSR1A)*, promoting the release of neuropeptides, such as neuropeptite Y and agouti-related protein, that enhance food intake [29,30].

Besides its effect on appetite, *GHRL* also influences peripheral metabolic processes such as fat mobilization and storage [31–35]. Recently, it was shown that UnG stimulates fatty acid oxidation and usage, while AcG induces deposition of lipids via lipogenesis in liver and adipose tissue [36,37]. This regulation is dependent on nutrient availability as *MBOAT4* responds to specific dietary lipids [27].

From an evolutionary standpoint, *GHRL* and *MBOAT4* have been found in mammals, birds, amphibians and fish [46] However, signs of sequence erosion were found in monotremes [47], a likely consequence of stomach loss (e. g. [48]). In turtles, initial studies were contradictory about the presence of the gene [49,50]. Yet, exhaustive sequence and expression analysis demonstrated the presence of a functional open reading frame in lizards, crocodilians and turtles [51–53]. By investigating the evolution of the hormone-enzyme duo in a larger genome dataset representative of all extant reptile lineages, including previously unexplored Serpentes and Chamaeleonidae, we deduce an unexpected pattern of gene loss [54], that may underscore the emergence of an extreme dietary phenotype.

## 2. Material and Methods

### (a) Datasets and retrieval of genomic regions corresponding to target orthologs

To establish the coding status of the target genes *GHRL* and *MBOAT4* in reptiles, we analysed an initial dataset consisting of 103 genomes of reptile species. These genomes were selected after applying a quality filter of 100 kilobase pairs (kbp) on scaffold N50 metric on the genomes available in GenBank in January 2024 (electronic supplementary material, file S1). tblastx from the blast package v2.13.0+ [55] was used to identify the presence and region of the target ortholog and target regions were extracted using samtools v1.15.1 [56] based on the top hit. As query, the *Gallus gallus* gene orthologs were used (*GHRL*: NM_001001131.2; *MBOAT4*: NM_001199289.2). For both datasets, genomic sequences obtained with tblastx hits on the reverse strand were reverse complemented using the tool “seq” from the seqtk package [57] with the “-r” flag. tblastx was also used to identify the presence and coordinates of the orthologs of the conserved flanking genes to establish if the local synteny is retained and the genomic regions between the nearest of these flanking orthologs were retrieved. We considered only tblastx hits in which the query and the subject frames were the same or in which the query frame was forward but the subject frame was reverse. Additional filters of 60% on percent identity and 20 bp of alignment length were applied. Whenever orthologs of *GHRL* or *MBOAT4* were not retrieved in downstream analyses due to phylogenetic distance to *Gallus gallus*, we repeated the analysis using the coding sequences of the orthologs in *Lacerta agilis* as query: (XM_033141137.1 (*LOC117041870/RPL32*); XM_033141136.1 (*LOC117041869/CAND2*); XM_033141139.1 (*SEC13*); XM_033141143.1 (*IRAK2*); XM_033141146.1 (*VHL*), XM_033141148.1 (*THUMPD3*).

### (b) Pseudogene and gene absence identification

Pseudochecker2.0 (command-line version of PseudoChecker [58,59] was used to detect premature stop codon (PSC) mutations, frameshift mutations, deletions of entire exons or genes, and mutations that disrupt splice sites. Based on the PseudoChecker outcome, we categorized genes as: pseudogenized, when there were disruptive mutations affecting more than 30% of the open reading frame; or absent, if the target ortholog could not be identified in a given genome but flanking orthologs were present in a contiguous region (*i.e.* no artefactual gaps). A threshold of 60% identity was established to determine if a specific exon was present. As functional coding references, we used the annotated orthologs in the *Trachemys scripta* genome (*GHRL*: XM_034776356.1; *MBOAT*4: XM_034771323.1) and, subsequently, in the *Lacerta agilis* (*GHRL*) and *Lacerta viridis* (*MBOAT4*) genome (using the exons and coding sequences predicted by PseudoChecker2) (electronic supplementary material, file S2). *Lacerta agilis* was not used as reference in MBOAT4 analyses as we detected a frameshift mutation in the MBOAT4 ortholog in this species. Lineages in which it was not possible to retrieve a functional ortholog due to missing exons were subjected to a following PseudoChecker2 analysis in an effort to retrieve the missing orthologous exons using a lower threshold (50%) for nucleotide percent identity. PseudoChecker2 was also used to create MACSE multiple sequence alignments for the three target genes (electronic supplementary material, file S3).

Exon alignments for the target genomic regions of *GHRL* and *MBOAT4* of representatives of certain groups (*Charina bottae*: Boidae; *Crotalus tigris*: Viperidae; *Furcifer pardalis*: Chamaeleonidae; *Liasis olivaceus*: Pythonidae; *Masticophis lateralis*: Colubridae; *Naja naja*: Elapidae; *Phrynocephalus forsythii*: Agamidae) were created in Geneious® 11.1.5 using the Map to Reference function with Highest Sensitivity and no Fine Tuning. Remaining settings were left on default. Disruptive exon mutations were then annotated from these alignments by hand, using a nucleotide identity cutoff of 50%. *Lacerta agilis* (*GHRL*: as predicted by PseudoChecker2), *Lacerta viridis* (*MBOAT4*: as predicted by PseudoChecker2), *Trachemys scripta* (*GHRL*: XM_034776356.1; *MBOAT*4: XM_034771323.1) and *Elgaria multicarinata* (*GHRL*: XM_063123404.1; *MBOAT4*: XM_063128442) were used as references. Disruptive exon mutations were manually annotated from these alignments. We further analyzed the genomes of 9 other species (electronic supplementary material, file S1) to confirm gene loss: three Pythonidae, one Boidae, three Agamidae, one Chamaeleonidae and one representative of blind snakes, Typhlopidae (the earliest branching lineage within Serpentes).

To demonstrate that the losses of *GHRL* were not due to artefactual gaps caused by low genome sequencing coverage, we extracted from the genome of the tiger rattlesnake, *Crotalus tigris* (GCF_016545835.1), the genomic region corresponding to the region starting from 2000 bp upstream of the *IRAK2* orthologue to 2000 bp downstream of the *SEC13* orthologue (as previously inferred by tblastx). Each region was then mapped using megablast [60] in the NCBI platform against the genomic read of its corresponding genome in the Sequence Read Archive (SRA) database [61]. The top 100 matching genomic reads were downloaded and mapped to the genomic region using minimap2 v.2.28 [62]. Samtools v1.9 [56] was used to convert the resultant SAM alignment into BAM format, as well as to sort and index the alignment in BAM format. The reads were then viewed in Integrative Genomics Viewer (IGV) (electronic supplementary material, figure S1) [63].

### (c) Alignment and phylogenetic analysis of GHRL orthologs

To demonstrate the conservation of *GHRL* across taxonomic groups, an alignment was created using Jalview [64] with protein sequences from representatives across Diapsida (birds and reptiles): NP_001001131.2 (*Gallus gallus*); XP_019392116.1 (*Crocodylus porosus*); XP_034632247.1 (*Trachemys scripta*); XP_032997031.1 (*Lacerta agilis*); CAOKZL010000534.1:19875-78509 (*Iguana delicatissima;* inferred from PseudoChecker2 results). To further assess if the detected orthologs are orthologs of *GHRL*, a Neighbor-joining tree [65] of the *GHRL* sequences retrieved by PseudoChecker2 and coding sequences from representative species across Tetrapoda was constructed using the MAFFT online service [66].

### (d) RNA-seq analysis

To assess if *GHRL* was expressed in snakes, the coding sequence of *Lacerta agilis* was used in a blast search (blastn) [55] against the stomach transcriptomic dataset of*Python molurus bivittatus* (SRA database under accessions SRX2182159 – fasting state, SRX2182169 – 1 day post feeding, SRX2182151 – 2 days post feeding and SRX2182156 – pool of all conditions) [67]. The obtained sequencing reads were then visualized in IGV to observe the splicing profiles of mature messenger RNA reads [63].

## 3. Results and Discussion

The genomic basis of the multiple adaptations observed in reptiles regarding feeding behavior and physiology remain poorly known. Here, we examined the genomic evolution of the *GHRL* and *MBOAT4* module across reptilian orders including members of the Squamata, Testudines, Crocodilia and Rhynchocephalia.

### (a) GHRL is absent or eroded in Serpentes and Chameleons

Using a combination of sequence comparisons and synteny analysis, we validated the presence of *GHRL* orthologues in all the examined species of lizards, turtles and tuatara (figure 2 and 3; electronic supplementary material, figure S2, files S4-S6). Phylogenetic analysis of the retrieved *GHRL* sequences yielded a topology displaying the expected relationships between Tetrapoda with well-supported clades, supporting the orthology of the gene sequences (electronic supplementary material, figure S3). We also found conserved synteny between the targets and their flanking genes (figure 2). The retrieved *GHRL* sequences display the typical features of the gene structure, as well as the characteristic third serine amino acid of the hormone peptide sequence (electronic supplementary material, figure S2), which is modified by the addition of an octanoyl group in the reaction catalysed by *MBOAT4* [27]. We merged the results from the two PseudoChecker2 analyses using as references the *GHRL* orthologs from *Trachemys scripta* and *Lacerta agilis* (a representative within Squamata) and found functional orthologs from across all species, except those within Serpentes (32 species), Chamaeleonidae (4 species), and the *Phrynocephalus* genus (Agamidae) (electronic supplementary material, files S3-S6). In the examined Colubridae, Elapidae and Viperidae species, no sequence with similarity to *GHRL* was found in the expected gene *locus* (figure 2). Homologous sequences of *GHRL* were found in *Liasis olivaceus* (Pythonidae) and *Charina bottae* (Boidae) but these were considered nonfunctional due to the presence of disruptive mutations (electronic supplementary material, file S6, figure 3). The mutational *GHRL* profile in the selected species shows a complex pattern (figure 3). For example, snakes from the Pythonidae and Boidae share premature stop codons (figure 3; electronic supplementary material, figure S4, file S6), including the first stop, which truncates the *GHRL* peptide after the 14^th^ amino acid (figure 3). In the Papuan ground boa, *Candoia aspera* (Boidae), we do not find this premature stop codon due to an upstream 4bp deletion at the end of the first exon (figure 3). There was also a 1bp deletion shared between *Charina bottae* and *Candoia aspera* (Boidae) in the 10^th^ codon, 4 codons before the start of the amino acid sequence that corresponds to the *Ghrl* peptide (figure 3). To further validate the loss of *GHRL*, we investigated the stomach transcriptome from the fasted *Python molurus bivittatus* (SRX2182159). In accordance, we found only two sequencing reads that aligned to *GHRL* (electronic supplementary material, file S7), despite clear expression of fasting-related genes [67]. The other stomach transcriptomic datasets only had 13 reads that aligned to *GHRL* (electronic supplementary material, file S8). Furthermore, alignment of these sequencing reads evidenced that these correspond to exon-intron (unspliced) reads (electronic supplementary material, figures S5-S6). Finally, exons retrieved through PseudoChecker for the other families within Serpentes with a lower identity threshold (50%) are unlikely to be real homologous sequences as evidenced by the MSA alignments (electronic supplementary material, file S3). Mapping of exons in Geneious supports this level of erosion/absence of *GHRL* (electronic supplementary material, file S9).

**Figure 2.**
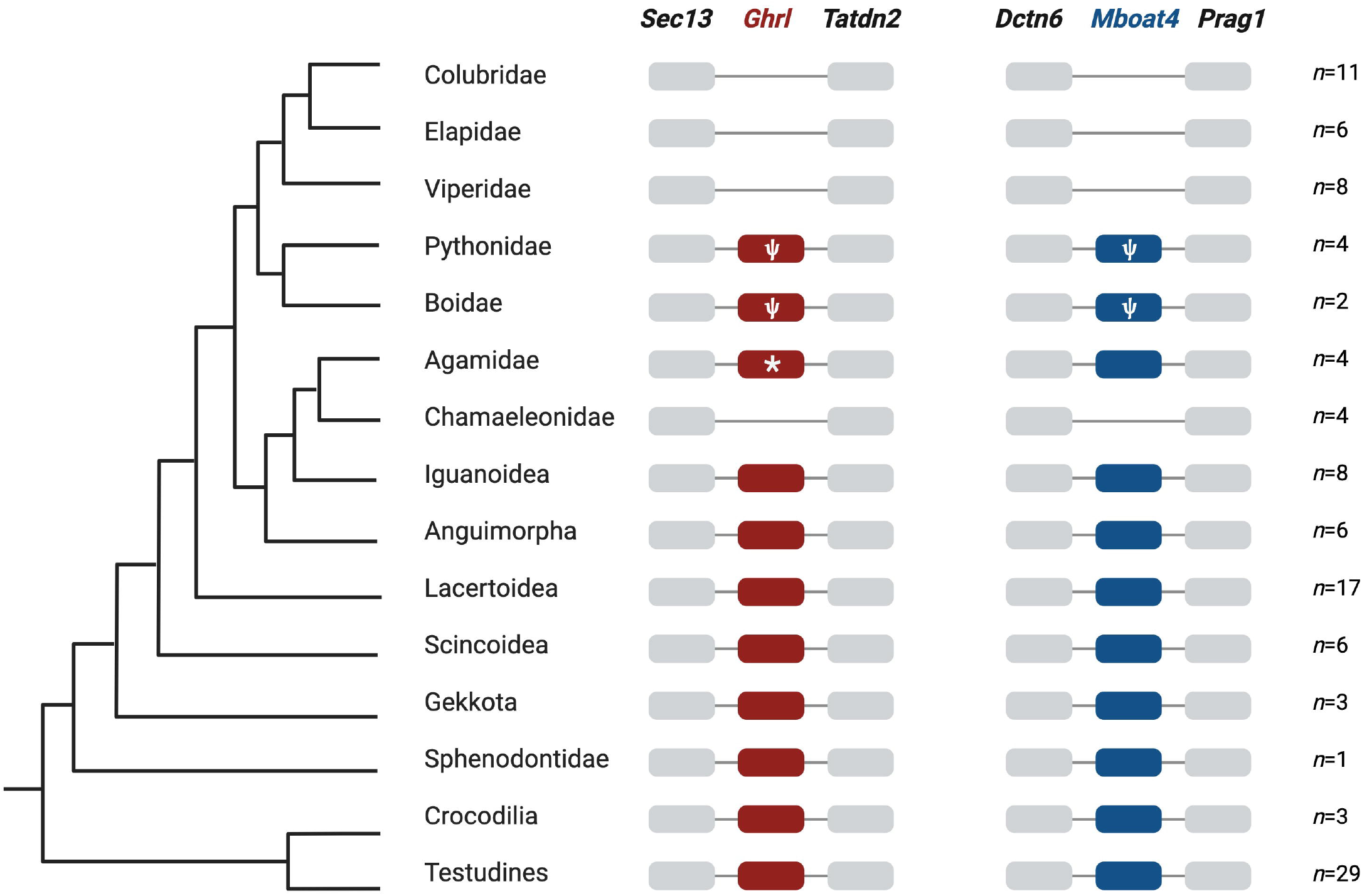
Absence of *GHRL* and *MBOAT4* in a locally conserved syntenic region across the reptile lineages (based on the phylogeny deduced by Pyron et al. *[68]*. Ghrl was considered present in a group if a seemingly functional ortholog was recovered in at least one of the species in that group.

**Figure 3.**
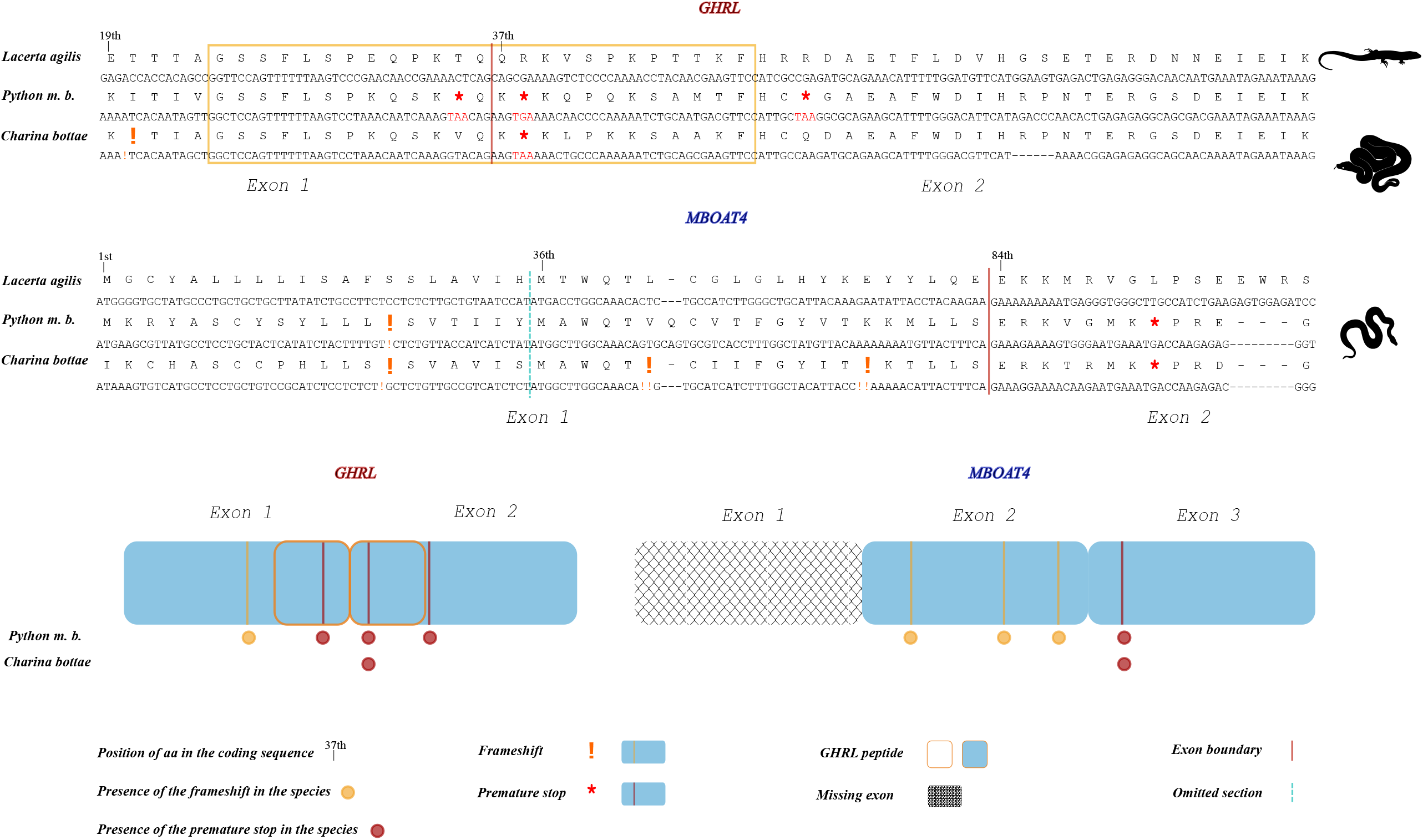
Mutational landscape in *GHRL* and *MBOAT4* across Serpentes selected species. (a) Alignment of *GHRL* (exons 1 and 2 shown here; peptide sequence highlighted in yellow) ortholog sequences in *Lacerta agilis, Python molurus bivittatus* and *Charina bottae* and of *MBOAT4* (exon 1 and start of exon 2 shown here, corresponding to exons 2 and 3 in Trachemys scripta) ortholog sequences in *Lacerta viridis, Python molurus bivittatus* and *Charina bottae*. (b) Presence and absence of exons (compared to the reference *Trachemys scripta*), as well as presence of deleterious mutations in *Python m. b.* and *Charina bottae.*

As for the blind snake, *Anilios bituberculatus*, representative of the earliest branching lineage within Serpentes, the high fragmentation in the target genomic regions made it impossible to extract and analyse. The *GHRL* genomic region was also highly fragmented in the genome of the veiled chameleon *Chamaeleo calyptratus* and we were unable to deduce any functional sequence from any of the Chamaeleonidae. Regarding Agamidae, even though no functional *GHRL* was found in the genus *Phrynocephalus* (represented by *P. forsythii* and *P. guinanensis*), we did detect a likely functional *GHRL* ortholog in *Intellagama lesueurii* and *Laudakia wui:* although the first codon of the second exon was first determined to be a premature stop codon, this is likely an artifact as that codon contains a possible splice site which had not been detected by the software, suggesting instead a deletion of the first codon (electronic supplementary material, file S4, figure S7). Thus, the loss of *GHRL* was likely not ancestral in this family, but the gene was instead lost independently in snakes, some agamids and chameleons.

Overall, we confirm an ancestral loss of *GHRL* in snakes and suggest independent loss events in chamaeleons and toadhead agamas (genus *Phrynocephalus*).

### (b) Sequence remnants of MBOAT4 are found in Serpentes genomes

Next, we investigated the coding status of the protein coded by the *MBOAT4* gene, an enzyme critical for the activation of the hormone *GHRL via* acylation (figure 1b). The search for functional orthologs of *MBOAT4* with PseudoChecker2 using the *Trachemys scripta* reference did not yield sequences homologous to *MBOAT4* across most of the Serpentes suborder, including species of Colubridae, Elapidae and Viperidae (figure 2). Yet, using the *MBOAT4* ortholog from *Lacerta viridis* as reference, we found sequences homologous to the first exon in representatives of the Pythonidae and Boidae, *Liasis olivaceus and Charina bottae* (figure 3). There is also a deletion of the 16^th^ nucleotide shared between the two families that shifts the frame of most of this exon (figure 3; electronic supplementary material, figure S8). However, in all species analyzed from these families, apart from *Charina bottae*, this frameshift is compensated by insertions at the end of this exon. We further found highly eroded sequences homologous to the second exon within *Pythonidae* and Boidae, as well as in other families within Serpentes (Elapidae and Viperidae). Besides the overall low degree of conservation of the amino acid sequences, there were also several conserved disruptive mutations (electronic supplementary material, file S6). *P. guinanensis* showed a loss of the second exon and a third exon with frameshift mutations and loss of the original 5’ splice site (electronic supplementary material, figure S9). Overall, we also found a high level of erosion of the *MBOAT4* gene and established the definitive loss of the *GHRL/MBOAT4* duo in snakes and chameleons.

### (c) Loss of the GHRL and MBOAT4 duo parallel specialized feeding strategies

We describe here the persistent erosion/deletion of the GHRL system in all snake lineages. Interestingly, snakes have specialized on the predation and digestion of very large prey relative to their own body mass, and this was achieved through specific anatomical and physiological adaptations. Concomitantly, snakes can tolerate extreme periods of fasting, lasting over a year between meals, despite taking less than 15 days to digest a large meal [69,70]. Thus, our findings of erosion/deletion of the *GHRL* system and the appetite stimulating effect of the hormone are consistent with the intermittent feeding done by snakes.

In fact, their ability to process such large meals after months of fasting suggests a drastic remodelling of the metabolic cues between the fed and the fasting states. In agreement, snakes also present a dramatic increase in the metabolic rate in response to digestion (the SDA response), reaching levels that are manyfold the standard metabolic rate, surpassing any other ectothermic sauropsid [13,14,21,71]. This response, which lasts for several days, is entirely supported by aerobic metabolism (as reviewed by Wang et al. [72]). In a fasting state, snakes are well adapted to save as much energy as possible and will, specifically, remain quiescent for long periods of time, thus saving energy in the skeletal muscle: especially ambush snakes, which do not employ active foraging [73–75]. The loss of *GHRL* suggests a lower need to stimulate fatty acid oxidation during fasting in the locomotor muscles of snakes compared to other groups of reptiles (figure 4). In addition, snake predation requires a burst of energy, best provided by glycolytic muscle fibers, which generate high-tension and enable faster movements [73,76]. In fact, reptiles sustain short-term increases in metabolic demands through glycolysis [77,78], efficiently supporting the physical effort required for predation [73,79,80]. As such, UnG may not be required to stimulate fatty acid oxidation in the skeletal muscle during fasting [81], providing a plausible trigger for the loss in this group. We suggest that the loss of the *GHRL* system evolved in parallel with the pattern of intermittent feeding of large meals observed in snakes. Interestingly, *GHRL* erosion in snakes is also detected in platypus and Eupasserines [47,82]. While, in monotremes, this event is linked to a radical shift in digestive physiology and anatomy remodelling (stomach loss) [48], in passerines, this genomic condition appears to be linked to fat deposition cycles [82].

**Figure 4.**
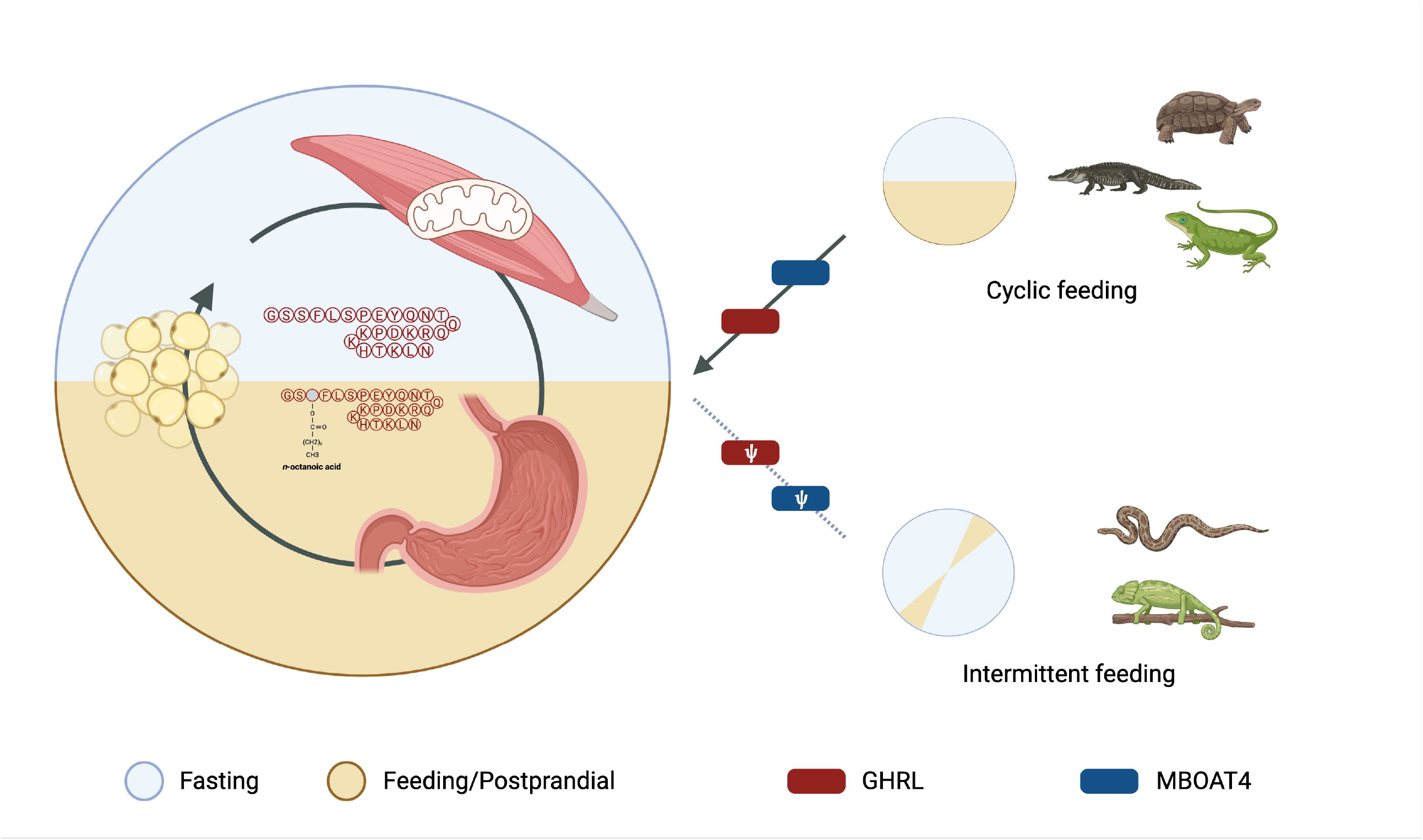
The loss of *GHRL*, which implicates a lack or reduction of fatty acid oxidation in the locomotory muscles, is lost due to the evolution of habits of intermittent feeding.

In addition to snakes, we also provide evidence for the loss of *GHRL* in the genus *Phrynocephalus* of the Agamidae family and chameleons. In the former, this type of foraging behavior is also observed, remaining motionless for long periods of time and moving through short bursts of rapid locomotion [83,84]. In addition, genetic, metabolic and morphological adaptations to deal with extreme environmental conditions are also observed in [85–91]. Finally, chameleons have been regarded as sit-and-wait foragers [92,93] or cruise foragers [94,95], a form of foraging characterized by scanning the environment for prey interspersed with short periods of locomotion [96]. In any case, they are ambush predators, supporting a similar behaviour-dependent metabolic scenario as that described in snakes.

## Conclusion

In conclusion, we uncovered the loss of the gene duo *GHRL* and *MBOAT4* in snakes, chamaeleons and toadhead agamas. This loss is a likely consequence of one of the many extreme physiological and behavioural adaptations that reptiles, particularly snakes, have evolved to survive in environments in which prey availability is uncertain and the need for energy budgeting is a constant. This study suggests that the intermittent feeding of these groups was accompanied by the loss of genes involved in energy homeostasis. The observed gene losses may reflect regressive evolution or an adaptive response with adaptive outcomes [54,97].

