## Supplemental Figures for "Ghrelin and Mboat4 are lost in Serpentes"

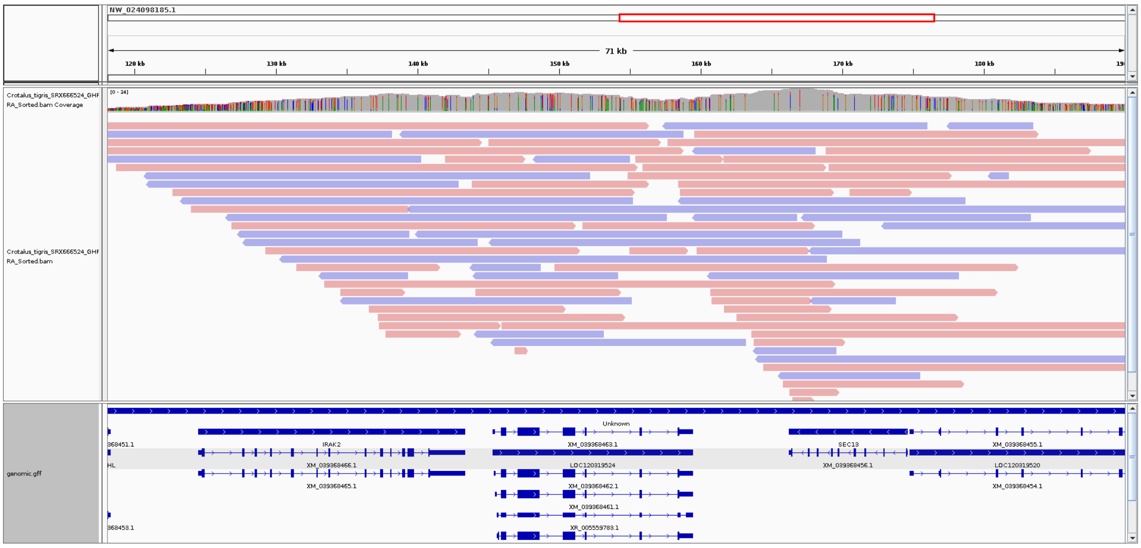


*Figure S1 – Reads mapped to the C. tigris genome demonstrating adequate coverage of the syntenic region of GHRL.*

| | |
|---|---|
| *Gallus gallus* | G S S F L S P T Y K N I Q Q Q K D T R K P T A R L H |
| *Crocodylus porosus* | G S S F L S P E Y P R T Q Q Q K D A K K P T T K L H |
| *Trachemys scripta* | G S S F L S P E Y Q N T Q Q R K D P K K - H T K L N |
| *Lacerta agilis* | G S S F L S P E Q P K T Q Q R K V S P K P T T K F H |
| *Iguana delicatissima* | G S S F L S P E Q P K - M Q R K V L Q K T P T K F H |

*Figure S2 - Alignment of Ghrl peptide sequences in representative species demonstrating conservation across diapsid lineages*

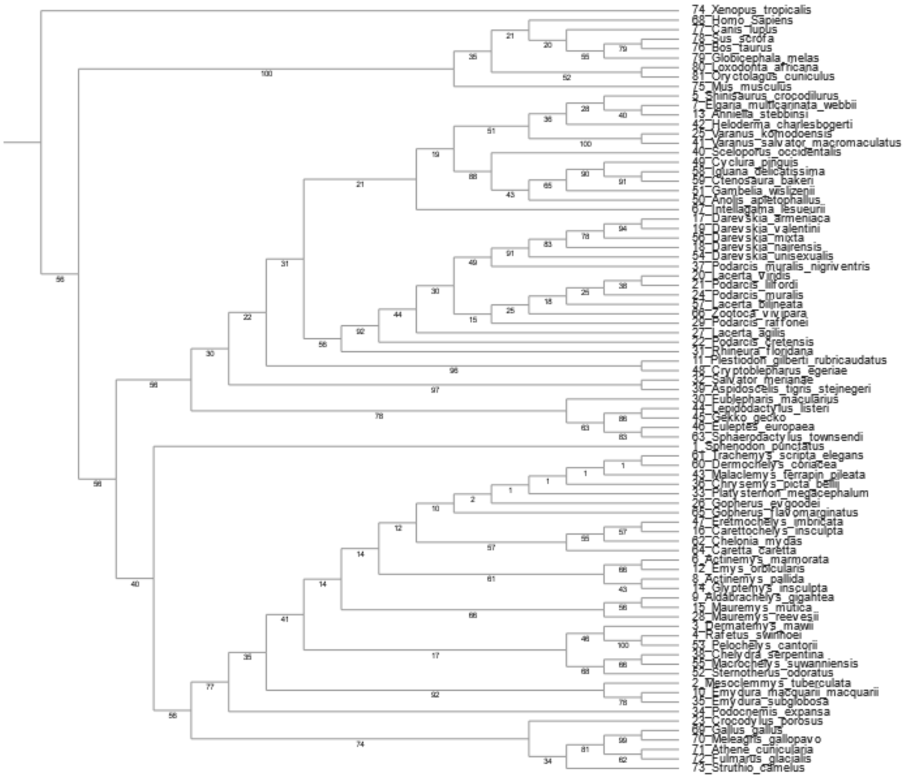


*Figure S3*- A Neighbor-joining phylogenetic analysis of the *Ghrl* sequences retrieved by PseudoChecker2 and representative species across Tetrapoda.

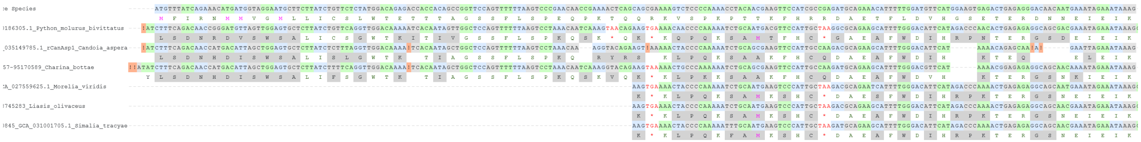

Figure S4 – MACSE multiple sequence alignment of sequences homologous to exons 1 and 2 of the *Ghrl* ortholog in *L. agilis* retrieved from the analysed genomes of the Pythonidae and Boidae families.

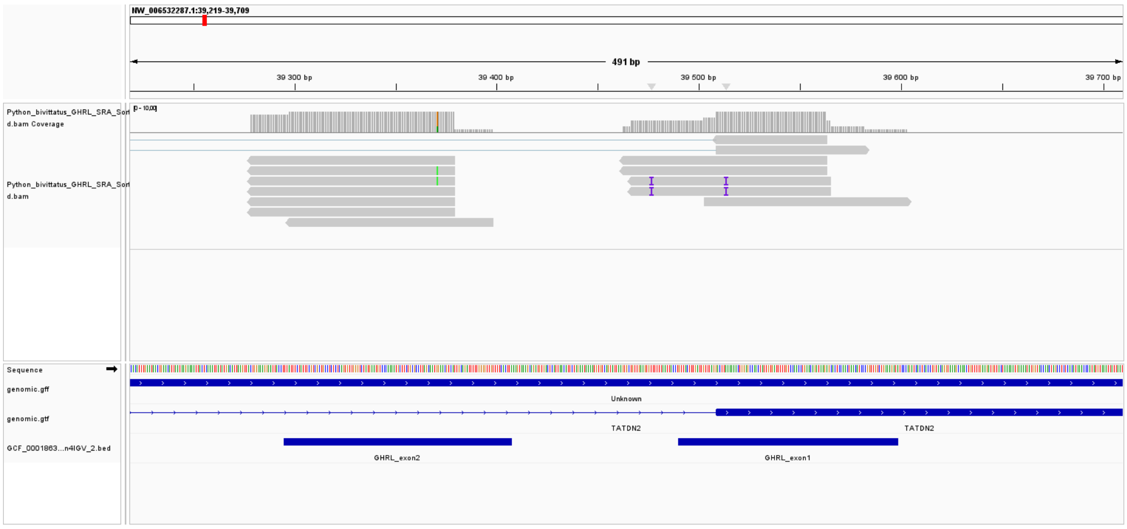


Figure S5 – Reads obtained from a blastn search of the *Ghrl* coding sequence (*L. agilis*) in the stomach transcriptomic datasets of *Python molurus bivittatus.*

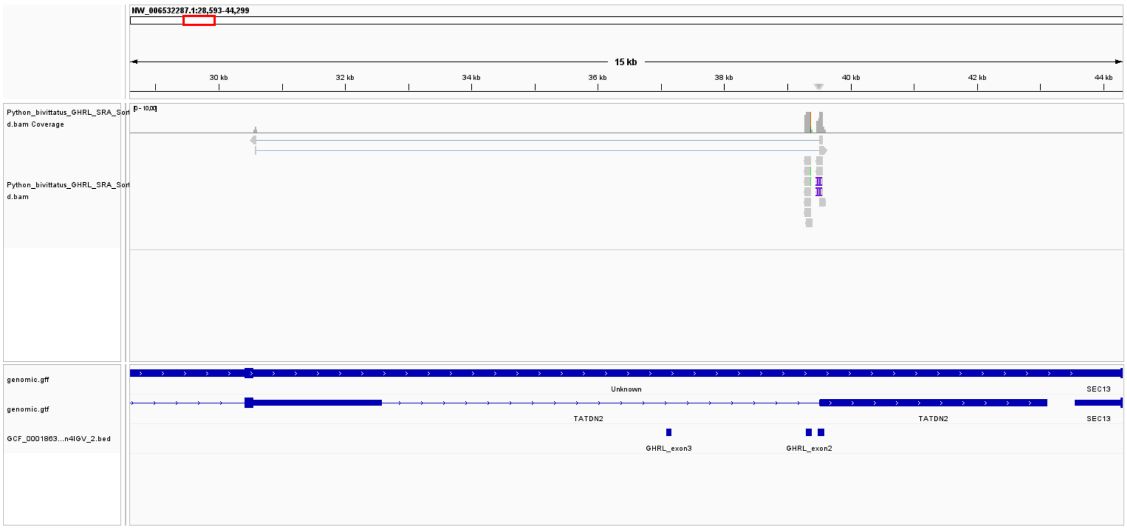

Figure S6 – Reads obtained from a blastn search of the *Ghrl* coding sequence (*L. agilis*) in the stomach transcriptomic datasets of *Python molurus bivittatus (zoomed out).*

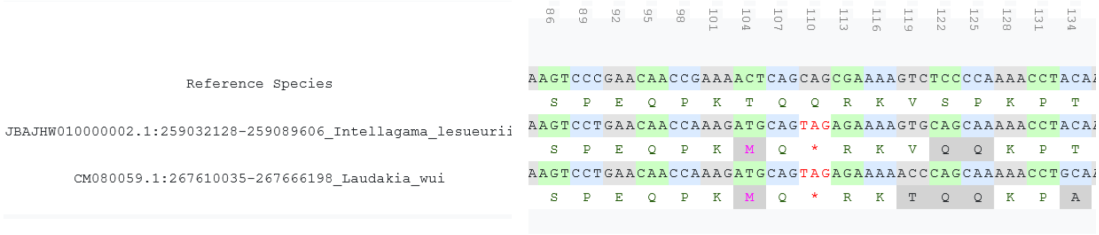


Figure S7 – MACSE alignment demonstrating the existence of a start codon in exon 2 of the MBOAT4 ortholog in *L. viridis*.

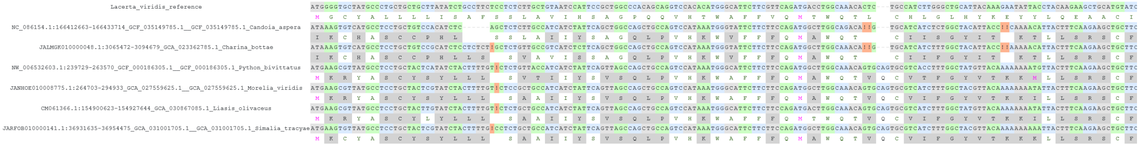

Figure S8 – MACSE multiple sequence alignment of sequences homologous to exons 1 and 2 of the *Mboat4* ortholog in *L. agilis* retrieved from the analysed genomes of the Pythonidae and Boidae families.

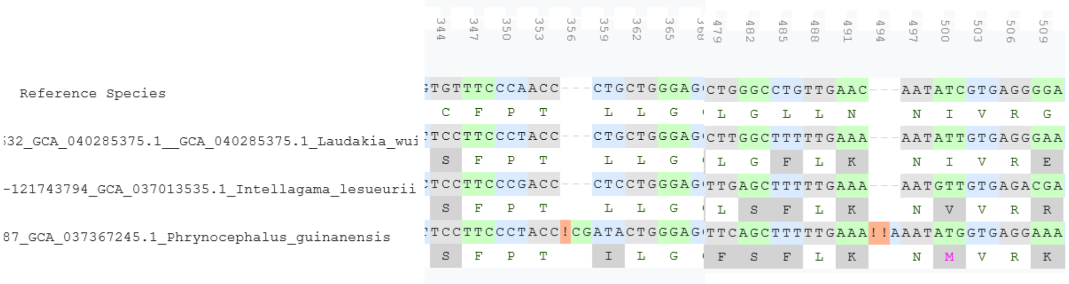

Figure S9 – MACSE multiple sequence alignment of sequences homologous to exons 2 and 3 of the *Mboat4* ortholog in *L. agilis* retrieved from the analysed genomes of the Agamidae.
