## Supplemental File 2 for "Ghrelin and Mboat4 are lost in Serpentes"

**References used for PseudoChecker2 analyses**

***Trachemys scripta Ghrl***

>Exon_1

TTTTCAGGTTTGATATAGTCTATGCGCCCAAGACACTCATGTTTCTCAGAAGCACTATGCTGGGAATTCT

CCTTATCTGCATTCTCTGGACAGAGACTACTATGGCTGGATCTAGTTTTTTAAGCCCTGAATATCAAAAC

ACACAG

>Exon_2

CAACGAAAGGATCCAAAAAAGCATACAAAGTTAAACCGCCGGGCTGCAGAAGGATTTTTGGATGCAGATGCAAGGCAGGCAGAAGGAGACAACAATGAAATAGAGATTAAG

>Exon_3

ATAACAGAAGACCAGTACCAGGAGTATGGACAAGTGCTGGAGAAGATCCTAGAGGACATTCTTGCAGAGGATACTAAAG

>Exon_4

AAACCCGAAATTGGCATGAGTTGAAACATGAAGATGTTACAAACTGATTAGAGTACAACGAAGTTCCACT

CATTGGCTTGGATTGCTTTAAATCTTGCTTTACATTGTGCAACAGATGATAATAGCATGTATTAAAAGAC

AAATGCATGAAGCTGAACAGTTCGCAGTTTTAAAAAAAATGACATTTTGTTGTGCAACAGATCTTTCGAT

GAGGTTGCAAGTAGTATATGTATGATAGACAGATAACAGCAAGCAAGGTCAAACACTGAAATCTAAATTG

TTTTCTTACCTTTCAAAACTATGTGACTGAAGCTACCTGTTCTAAACAAGTGAGGGAAAAACTGTGTTAA

TTTACCATATGCTTCTTTCAGCAGAAAATAAAGCTAGTGTTGAAACCGC

>CDS

ATGTTTCTCAGAAGCACTATGCTGGGAATTCTCCTTATCTGCATTCTCTGGACAGAGACTACTATGGCTGGATCTAGTTTTTTAAGCCCGAATATCAAAACACACAGCAACGAAAGGATCCAAAAAAGCATACAAAGTTAAACCGCCGGGCTGCAGAAGGATTTTTGGATGCAGATGCAAGGCAGGCAGAAGGAGACAACAATGAAATAGAGATTAAGATAACAGAAGACCAGTACCAGGAGTATGGACAAGTGCTGGAGAAGATCCTAGAGGACATTCTTGCAGAGGATACTAAAGAAACCCGAAATTGGCATGAGTTGAAACATGAAGATGTTACAAACTGA

***Trachemys scripta Mboat4***

>Exon_1

ATGAAGGCAGCAGTCACGGGTTATAATCTGAGGAAAGAGTGGGGAGAGGAAATCAATGCACCAAGTTCACTCAGGTCTTCTGGAGTTAACGGGACTTGCAG

>Exon_2

GTACGCCTTGCTCCTGGTGGGAGGCTGCCTTCTCGCGTGTGCTGCCATGGGATGGTATGCCCTGCTGCTC

TTCATCCCCACCTTCTGCTCCGTGGCCATTTTCCACTCAGTCGGCCCACTGCAGGTCCACACATGGGCAT

TCGTGCTCCAGATGTCCTGGCAGACTCTCTGCCACCTGGGCCTGCACTACAGAGAACATTATCTGCAAGG

AGCCCCGTGCATCAG

>Exon_3

GTTGACAATCGCCCTCTCGTCGCTCATGCTGCAGACGCAGAAGGTCACGTCGCTGGCTCTGGATATCCAC

GAAGGGAAGGTGACTATGGCAGCTGAGTGGGGGGATGGGAGGGAGCCACTGCTGCGGGCGCTGCCTCTATGCAGCTACCTGCTCTTTTTCCCTGCCCTGCTGGGAGGCCCGCTATGCTCCTTTCGGAGATTTCAGGACCAGATCCAGGGCTCATGCGCTTCACGCCCCACACCTCCCTGGAAGGCTGCAGGCCAGAAATGCCTTCGTGCCCTGGCCCTGCACCTGCTGAGAACGGCTGTGAGGAGCTGCGTGGCCCCGCTGGCCCGCATGACGGACTGCACCGGCTTTGGCTGCGTGTTTGTCATGTGGCGTTCCGCCCTGCTCTTCAAACTGGCTTATTACTCCCAGTGGGTGCTCGACGAGGCTCTCCTCAGCGCAGCGGGCTTTGGGCTGGAGCTTGGCCACGCCCCCGGTGCAGAGGCTGCCTGCGGCGACCTCTCTGATGCAGACATATGGACCTTGGAGACCACCAACAGAATAGCCCTCTTCACCAGGACCTGGAACAAGAGCACTTCCCGGTGGCTGAGGAGACTTGTCTTCCAGCGCAGCCCGGCCCAGCCCCTCTTGGCCACCTTTGCCTTCTCGGCCTGGTGGCACGGGCTCCACCCAGGGCAAGTGTTTGGCTTCTTGTGCTGGGCAGCGATGGTGGAGGCCGATTACCGGATTCACCCTTTTCTCCGTTCACTTGCAAAGTCCTGGCACACCAAGGTGCTGTATCAGGCCCTGACCTGGGTCCAGACCCAGCTGATCATTGCGTACATCACGGTTGCGGTGGAGATGAGGAGCTTCTCTGCGCTCTGGCTGCTGGGCGCCTCCTACAACAGCTTCTTCCCTCTCCTGTACGGTGTTTCACTGCTGTGGCTGGTCGCGAGAGCCAAAGAAAAATGTGTCTGACCCACACTATTCTCCCTCCCACACACACACTATTCACCAGGAGCCATAGAGCGTAGGAGAGATGGAGATTGCCCCCTTGGTAGGGAGAGGTTTTCTCTAGGTCTAGGTATGCCCCTAGTCCCCCCTAAACCCATCATCTCATCACATAAGATAGACTCTCACATTCTAAGGCCAGGAGGGACCATTAGATCATCCAGTCTGACCTGCTCCATAGCACACACCAGAGAACCTCACCCAGTGATTCCCACATCCGGCAAATAACCCAAGGTTGGATCAGAGTGTATCTTTTAGAAGAAGACATCCTGTCTTGCTTAAAGGGTCCAAGTGATGAAGAATTTATCATAGAA

>CDS

ATGAAGGCAGCAGTCACGGGTTATAATCTGAGGAAAGAGTGGGGAGAGGAAATCAATGCACCAAGTTCACTCAGGTCTTCTGGAGTTAACGGGACTTGCAGGTACGCCTTGCTCCTGGTGGGAGGCTGCCTTCTCGCGTGTGCTGCCATGGGATGGTATGCCCTGCTGCTCTTCATCCCCACCTTCTGCTCCGTGGCCATTTTCCACTCAGTCGGCCCACTGCAGGTCCACACATGGGCATTCGTGCTCCAGATGTCCTGGCAGACTCTCTGCCACCTGGGCCTGCACTACAGAGAACATTATCTGCAAGGAGCCCCGTGCATCAGGTTGACAATCGCCCTCTCGTCGCTCATGCTGCAGACGCAGAAGGTCACGTCGCTGGCTCTGGATATCCACGAAGGGAAGGTGACTATGGCAGCTGAGTGGGGGGATGGGAGGGAGCCACTGCTGCGGGCGCTGCCTCTATGCAGCTACCTGCTCTTTTTCCCTGCCCTGCTGGGAGGCCCGCTATGCTCCTTTCGGAGATTTCAGGACCAGATCCAGGGCTCATGCGCTTCACGCCCCACACCTCCCTGGAAGGCTGCAGGCCAGAAATGCCTTCGTGCCCTGGCCCTGCACCTGCTGAGAACGGCTGTGAGGAGCTGCGTGGCCCCGCTGGCCCGCATGACGGACTGCACCGGCTTTGGCTGCGTGTTTGTCATGTGGCGTTCCGCCCTGCTCTTCAAACTGGCTTATTACTCCCAGTGGGTGCTCGACGAGGCTCTCCTCAGCGCAGCGGGCTTTGGGCTGGAGCTTGGCCACGCCCCCGGTGCAGAGGCTGCCTGCGGCGACCTCTCTGATGCAGACATATGGACCTTGGAGACCACCAACAGAATAGCCCTCTTCACCAGGACCTGGAACAAGAGCACTTCCCGGTGGCTGAGGAGACTTGTCTTCCAGCGCAGCCCGGCCCAGCCCCTCTTGGCCACCTTTGCCTTCTCGGCCTGGTGGCACGGGCTCCACCCAGGGCAAGTGTTTGGCTTCTTGTGCTGGGCAGCGATGGTGGAGGCCGATTACCGGATTCACCCTTTTCTCCGTTCACTTGCAAAGTCCTGGCACACCAAGGTGCTGTATCAGGCCCTGACCTGGGTCCAGACCCAGCTGATCATTGCGTACATCACGGTTGCGGTGGAGATGAGGAGCTTCTCTGCGCTCTGGCTGCTGGGCGCCTCCTACAACAGCTTCTTCCCTCTCCTGTACGGTGTTTCACTGCTGTGGCTGGTCGCGAGAGCCAAAGAAAAATGTGTCTGA

***Lacerta agilis Ghrl***

>Exon_1

ATGTTTATCAGAAACATGATGGTAGGAATGCTTCTTATCTGTTCTCTATGGACAGAGACCACCACAGCCGGTTCCAGTTTTTTAAGTCCCGAACAACCGAAAACTCAG

>Exon_2

CAGCGAAAAGTCTCCCCAAAACCTACAACGAAGTTCCATCGCCGAGATGCAGAAACATTTTTGGATGTTCATGGAAGTGAGACTGAGAGGGACAACAATGAAATAGAAATAAAG

>Exon_3

ATCTCAGAAGATCAGTACAAAGACTATGGCCCAATACTGGAGAAGCTTCTGGAAGACATACTTGTTGAAGGCAATAAAG

>CDS

ATGTTTATCAGAAACATGATGGTAGGAATGCTTCTTATCTGTTCTCTATGGACAGAGACCACCACAGCCGGTTCCAGTTTTTTAAGTCCCGAACAACCGAAAACTCAGCAGCGAAAAGTCTCCCCAAAACCTACAACGAAGTTCCATCGCCGAGATGCAGAAACATTTTTGGATGTTCATGGAAGTGAGACTGAGAGGGACAACAATGAATAGAAATAAAGATCTCAGAAGATCAGTACAAAGACTATGGCCCAATACTGGAGAAGCTTCTGGAAGACATACTTGTTGAAGGCAATAAAG

***Lacerta viridis Mboat4***

>Exon_1

GTACACTTTCCTCCTGGTGGGTGGTTTCCTCCTGGCCTGGGCTGCCATGGGGTGCTATGCCCTGCTGCTGCTTATATCTGCCTTCTCCTCTCTTGCTGTAATCCATTCCGCTGGCCCACAGCAGGTCCACACATGGGCATTCTTCGTTCAGATGACCTGGCAAACACTCTGCCATCTTGGGCTGCATTACAAAGAATATTACCTACAAGAAGCTGCATGTATCAG

>Exon_2

ATTGCCCATCGCCCTTTCTGCCCTCATGCTGATGACCCAGAAAGTCACCTCACTGGCTTTGGATATCCATGAAAAAAAAATGAGGGTGGGCTTGCCATCTGAAGAGTGGAGATCCTTTTGTTGGCACCTGCTCCAGGCACTGCCACTGTGTACTTATCTGCTGTGTTTCCCAACCCTGCTGGGAGGCCCTCTGTATTCTTTCCGCAGATTTCAGGCATGGGTAAGGTATTCCAAGGCTCCGTTTTCTTCGGGTCTCCTCTGGGCTGCCACTCGAAAAGGCCTGGGGGCTCTGACTCTGGGCCTGTTGAACAATATCGTGAGGGGATACATTTCCCCTCTGGACGACCTAATCGACTGCACCCACTTTGACTGCGTTTACATCATGTGGACCTCAGCCCTGTCCTTCAGGCTCACCTATTACTCTCACTGGCTGCTTGACGAGTCCCTCTTCCTTGCTGCTGGTTTGGGGCTGGATCTTGGCCACCACCAATATTCAGCAGCTGCCGACAGAGTTGTCATGGACACAGACATTTGGACCCTGGAAACAACCAACACAATTGCTGGCTTTACCCGAACATGGAACAAGAGCACAGCCCAGTGGCTGAGGCGGCTCATATTCCAGCAGAGTAGTTCACATCCTCTCCTGGCCACCTTTGCCTTCTCAGCCTGGTGGCATGGCCTCCACCCTAGCCAGGTCTTTGGGTTTTTGTGCTGGGCCGTTATGGTGGAAGCAGACTACCGCTTCCATCGCTTTTTTGGTTCAGTGGCAAAATCCCGGCTTCAGAAGCTGCTGTACCAAACTGTGACCTGGTGCCATACACAACTGGTGGTGGCGTACATTATGATTGCTGTTGAGATTAGGAGTGTGTCCATGCTTTGGCAGCTGTTGTCTTCGTACAATAGCTTCTTTCCACTAGTCTCTGTCACTGTGCTCCTCCTGTTAGTAAAGAAATGAGTGAGC

>CDS

ATGGGGTGCTATGCCCTGCTGCTGCTTATATCTGCCTTCTCCTCTCTTGCTGTAATCCATTCCGCTGGCCCACAGCAGGTCCACACATGGGCATTCTTCGTTCAGATGACCTGGCAAACACTCTGCCATCTTGGGCTGCATTACAAAGAATATTACCTACAAGAAGCTGCATGTATCAGATTGCCCATCGCCCTTTCTGCCCTCATGCTGATGACCCAGAAAGTCACCTCACTGGCTTTGGATATCCATGAAAAAAAAATGAGGGTGGGCTTGCCATCTGAAGAGTGGAGATCCTTTTGTTGGCACCTGCTCCAGGCACTGCCACTGTGTACTTATCTGCTGTGTTTCCCAACCCTGCTGGGAGGCCCTCTGTATTCTTTCCGCAGATTTCAGGCATGGGTAAGGTATTCCAAGGCTCCGTTTTCTTCGGGTCTCCTCTGGGCTGCCACTCGAAAAGGCCTGGGGGCTCTGACTCTGGGCCTGTTGAACAATATCGTGAGGGGATACATTTCCCCTCTGGACGACCTAATCGACTGCACCCACTTTGACTGCGTTTACATCATGTGGACCTCAGCCCTGTCCTTCAGGCTCACCTATTACTCTCACTGGCTGCTTGACGAGTCCCTCTTCCTTGCTGCTGGTTTGGGGCTGGATCTTGGCCACCACCAATATTCAGCAGCTGCCGACAGAGTTGTCATGGACACAGACATTTGGACCCTGGAAACAACCAACACAATTGCTGGCTTTACCCGAACATGGAACAAGAGCACAGCCCAGTGGCTGAGGCGGCTCATATTCCAGCAGAGTAGTTCACATCCTCTCCTGGCCACCTTTGCCTTCTCAGCCTGGTGGCATGGCCTCCACCCTAGCCAGGTCTTTGGGTTTTTGTGCTGGGCCGTTATGGTGGAAGCAGACTACCGCTTCCATCGCTTTTTTGGTTCAGTGGCAAAATCCCGGCTTCAGAAGCTGCTGTACCAAACTGTGACCTGGTGCCATACACAACTGGTGGTGGCGTACATTATGATTGCTGTTGAGATTAGGAGTGTGTCCATGCTTTGGCAGCTGTTGTCTTCGTACAATAGCTTCTTTCCACTAGTCTCTGTCACTGTGCTCCTCCTGTTAGTAAAGAAATGA
