## Supplemental File 7 for "Ghrelin and Mboat4 are lost in Serpentes"

>gnl|SRA|SRR4280484.656885.2 HWI-
ST132:497:C016LACXX:2:1104:14342:113219
CAACCCCAAAAATCTGCAATGACGTTCCATTGCTAAGGCGCAGAAGCATTTTGGGACATTCATAGACC
CA
ACACTGAGAGAGGCAGCGACGAAATAGAAAT
>gnl|SRA|SRR4280484.3580637.1 HWI-
ST132:497:C016LACXX:2:1303:12937:90064
GAGCTGAAGAGGAAATGCTTCTGTAAAGACTTAAAAAACTGGAGCCAACTATTGTGATTTTTGTCCAA
CC
TGAACAGATAAGAGCACTCCAACTAACATCC
