## Supplemental File 8 for "Ghrelin and Mboat4 are lost in Serpentes"

>gnl|SRA|SRR4280494.101450.1 HWI-
ST132:497:C016LACXX:2:1101:1188:124431
GTTTAGGACTTAAAAAACTGGAGCCAACTATTGTGATTTTTGTCCAACCTGAACAGATAAGAGCACTC
CA
ACTAACATCCCGGTTGTCTGAAAGATATCCC
>gnl|SRA|SRR4280476.10134147.2 HWI-
ST132:497:C016LACXX:2:2206:12340:110787
CTTATCTGTTCAGGTTGGACAAAAATCACAATAGTTGGCTCCAGT
TTTTTAAGTCAGAAGCATTTCCTCTTCAGCTCTTCCCCTTCTCTTTTGCAGCTTCT
>gnl|SRA|SRR4280481.9228597.1 HWI-
ST132:497:C016LACXX:3:2207:6712:51278
TGACGTTCCATTGCTAAGGCGC
AGAAGCATTTTGGGACATTCATAGACCCAACACTGAGAGAGGCAGCGACGAAATAGAAATAAAGGTAA
TTCCTTTCCAG-
>gnl|SRA|SRR4280481.5552750.1 HWI-
ST132:497:C016LACXX:3:1308:3837:62603
TGACGTTCCATTGCTAAGGCGC
AGAAGCATTTTGGGACATTCATAGACCCAACACTGAGAGAGGCAGCGACGAAATAGAAATAAAGGTAA
TTCCTTTCCAG-
>gnl|SRA|SRR4280481.5548088.1 HWI-
ST132:497:C016LACXX:3:1308:13155:59054
TGACGTTCTATTGCTAAGGCGC
AGAAGCATTTTGGGACATTCATAGACCCAACACTGAGAGAGGCAGCGACGAAATAGAAATAAAGGTAA
TTCCTTTCCAG-
>gnl|SRA|SRR4280481.10441705.2 HWI-
ST132:497:C016LACXX:3:2304:8050:73962
CTTATCTGTTCAGGTTGGACAAAAATCACAATAGTTGGCTCCAGT
TTTTTAAGTCCTAAACAATCAAAGTAACAGGTAATTAAAAAACACTTCAACAGCAC
>gnl|SRA|SRR4280481.1461899.2 HWI-
ST132:497:C016LACXX:3:1107:20548:64141
CTCTTATCTGTTCAGGTTGGACAAAAATCACAATAGTTGGCTCCAGT
TTTTTTAAGTCCTAAACAATCAAAGTAACAGGTAATTAAAAAAACACTTCAACA
>gnl|SRA|SRR4280481.9228597.1 HWI-
ST132:497:C016LACXX:3:2207:6712:51278
TGACGTTCCATTGCTAAGGCGCAGAAGCATTTTGGGACATTCATAGACCCAACACTGAGAGAGGCAGC
GA
CGAAATAGAAATAAAGGTAATTCCTTTCCAG
>gnl|SRA|SRR4280481.5552750.1 HWI-
ST132:497:C016LACXX:3:1308:3837:62603
TGACGTTCCATTGCTAAGGCGCAGAAGCATTTTGGGACATTCATAGACCCAACACTGAGAGAGGCAGC
GA
CGAAATAGAAATAAAGGTAATTCCTTTCCAG
>gnl|SRA|SRR4280481.5548088.1 HWI-
ST132:497:C016LACXX:3:1308:13155:59054
TGACGTTCTATTGCTAAGGCGCAGAAGCATTTTGGGACATTCATAGACCCAACACTGAGAGAGGCAGC
GA
CGAAATAGAAATAAAGGTAATTCCTTTCCAG
>gnl|SRA|SRR4280481.10441705.2 HWI-
ST132:497:C016LACXX:3:2304:8050:73962
CTTATCTGTTCAGGTTGGACAAAAATCACAATAGTTGGCTCCAGTTTTTTAAGTCCTAAACAATCAAA
GT
AACAGGTAATTAAAAAACACTTCAACAGCAC
>gnl|SRA|SRR4280481.1461899.2 HWI-
ST132:497:C016LACXX:3:1107:20548:64141

CTCTTATCTGTTCAGGTTGGACAAAAATCACAATAGTTGGCTCCAGTTTTTTTAAGTCCTAAACAATC
AA
AGTAACAGGTAATTAAAAAAACACTTCAACA
>gnl|SRA|SRR4280484.656885.2 HWI-
ST132:497:C016LACXX:2:1104:14342:113219
CAACCCCAAAAATCTGCAATGACGTTCCATTGCTAAGGCGCAGAAGCATTTTGGGACATTCATAGACC
CA
ACACTGAGAGAGGCAGCGACGAAATAGAAAT
>gnl|SRA|SRR4280484.3580637.1 HWI-
ST132:497:C016LACXX:2:1303:12937:90064
GAGCTGAAGAGGAAATGCTTCTGTAAAGACTTAAAAAACTGGAGCCAACTATTGTGATTTTTGTCCAA
CC
TGAACAGATAAGAGCACTCCAACTAACATCC
