## Supplementary figures and images for "Ghrelin and Mboat4 are lost in Serpentes"

### Supplemental File 9

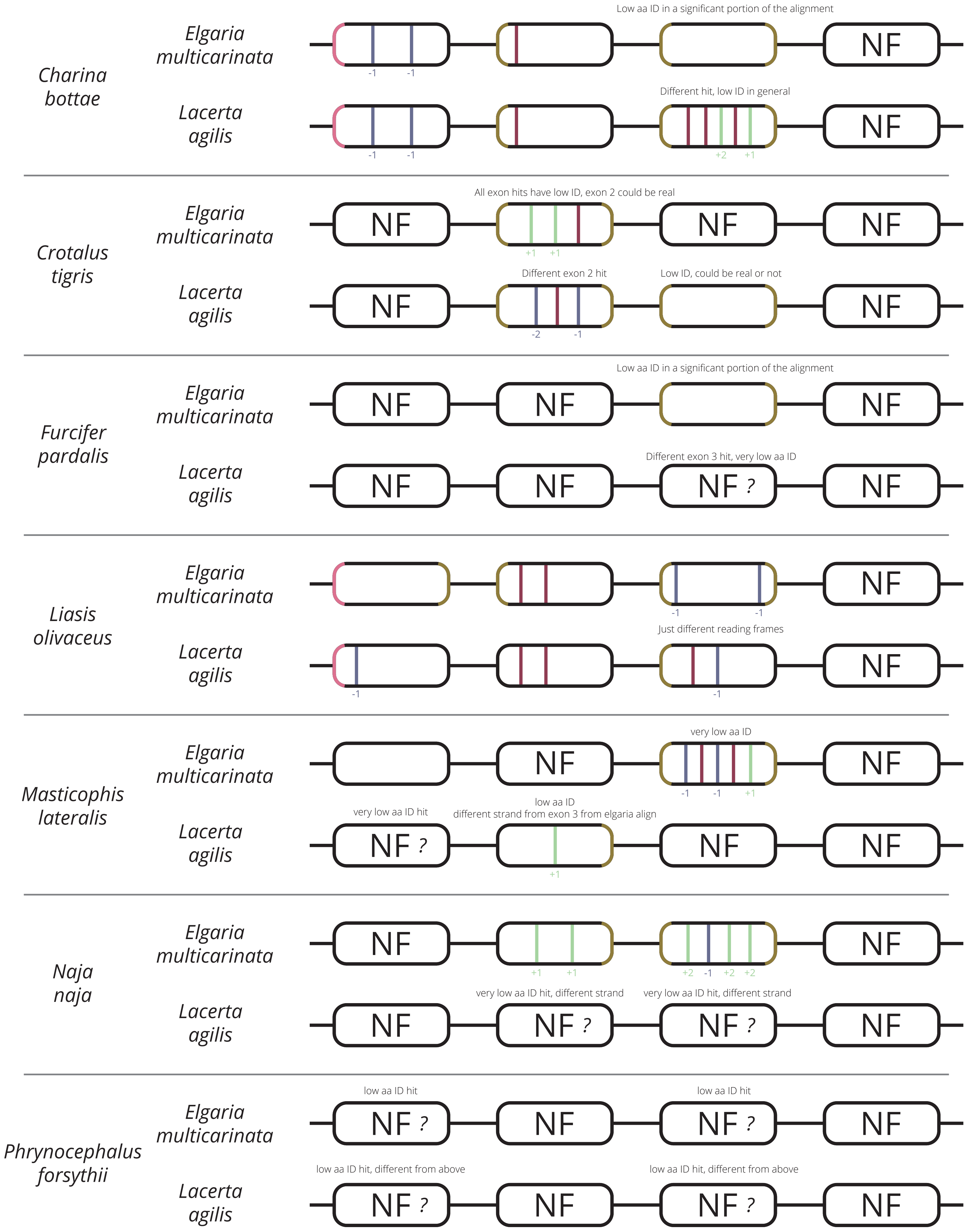

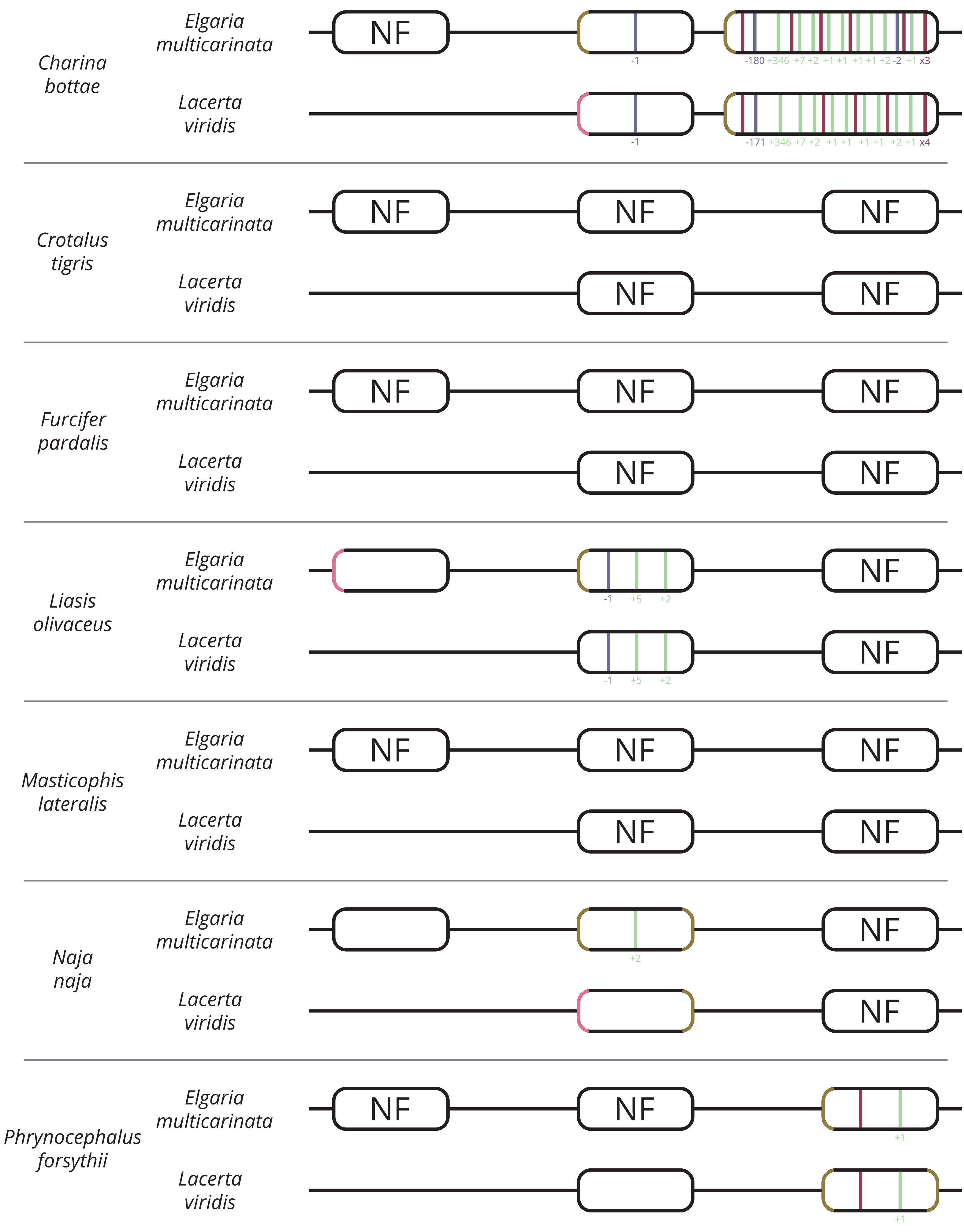
